# Shifts in carbon partitioning by photosynthetic activity increase terpenoid synthesis in glandular trichomes

**DOI:** 10.1101/2022.09.29.510054

**Authors:** Nima P. Saadat, Marvin van Aalst, Alejandro Brand, Oliver Ebenhöh, Alain Tissier, Anna B. Matuszyńska

**Affiliations:** Institute of Theoretical and Quantitative Biology, Heinrich Heine University Düsseldorf, 40225 Düsseldorf, Germany; Leibniz-Institut für Pflanzenbiochemie, Weinberg 3, 06120 Halle, Germany; Cluster of Excellence on Plant Sciences, Heinrich Heine University Düsseldorf, Universitätsstraße 1, 40225 Düsseldorf, Germany; Computational Life Science, Department of Biology, RWTH Aachen University, Worringerweg 1, 52074 Aachen, Germany

**Keywords:** bioenergetics, glandular trichomes, photosynthesis, stoichiometric model, secondary metabolites, terpenes

## Abstract

Several commercially important secondary metabolites are produced and accumulated in high amounts by glandular trichomes, giving the prospect of using them as metabolic cell factories. Due to extremely high metabolic fluxes through glandular trichomes, previous research focused on how such flows are achieved. The question regarding their bioenergetics became even more interesting with the discovery of photosynthetic activity in some glandular trichomes. Despite recent advances, how primary metabolism contributes to the high metabolic fluxes in glandular trichomes is still not fully elucidated. Using computational methods and available multi-omics data, we first developed a quantitative framework to investigate the possible role of photosynthetic energy supply in terpenoid production and next tested experimentally the simulation-driven hypothesis. With this work, we provide the first reconstruction of specialised metabolism in Type-VI photosynthetic glandular trichomes of *Solanum lycopersicum*. Our model predicted that increasing light intensities results in a shift of carbon partitioning from catabolic to anabolic reactions driven by the energy availability of the cell. Moreover, we show the benefit of shifting between isoprenoid pathways under different light regimes, leading to a production of different classes of terpenes. Our computational predictions were confirmed *in vivo*, demonstrating a significant increase in production of monoterpenoids while the sesquiterpenes remained unchanged under higher light intensities. The outcomes of this research provide quantitative measures to assess the beneficial role of chloroplast in glandular trichomes for enhanced production of secondary metabolites and can guide the design of new experiments that aim at modulating terpenoid production.

**GRAPHICAL ABSTRACT:** 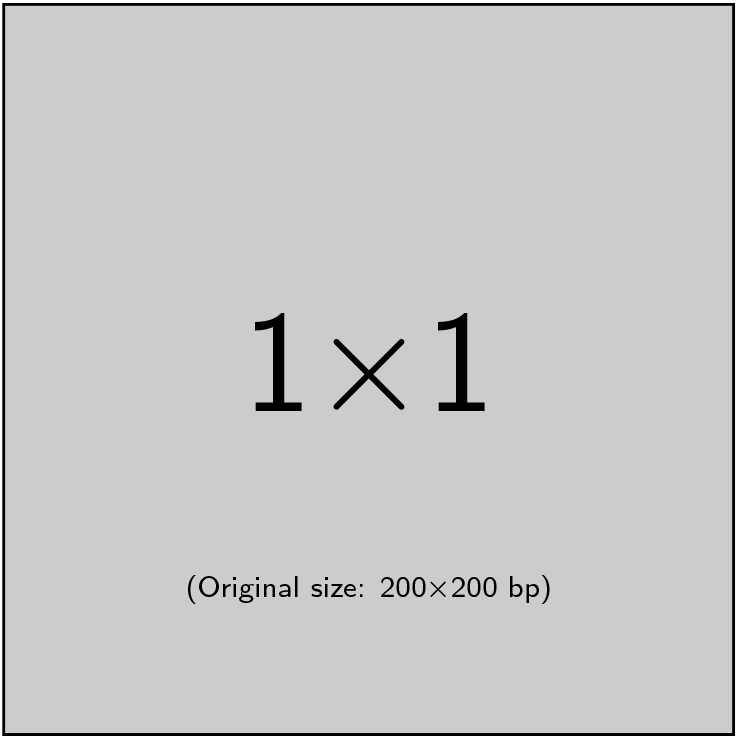

Please check the journal’s author guidelines for whether a graphical abstract, key points, new findings, or other items are required for display in the Table of Contents.

## 1 INTRODUCTION

Most plant species exhibit cellular outgrowths of their epidermis called trichomes. Due to their often species-specific characteristic, many criteria for classification exist, the most popular one being the division into non-glandular and **glandular trichomes** (GT) [1]. Whilst non-glandular trichomes serve more as a physical and mechanical defence against biotic and abiotic stresses, all GTs are characterised by the ability to synthesize and accumulate vast amounts of valuable specialised (secondary) metabolites. Due to extremely high metabolic fluxes in these organs, production of some metabolites can reach up to 20% of the leaf dry weight [2], GTs are often referred to as true **metabolic cell factories** [3]. Products of GTs include terpenoids, phenylpropanoids, flavonoids, fatty acid derivatives, and acyl sugars [4] exhibiting antifungal, insecticide, or pesticide properties. Thereby GTs are not only incredibly important to plant fitness, as they contribute to the chemical arsenal of plants, but are also of relevance to multiple industries.

The key carbon source in most GTs of tomatoes is sucrose which is converted into a multitude of organism-specific metabolites in the glands [5]. The massive productivity of hydrocarbon compounds implies, however, a supply of adequate amounts of not only carbon, energy, and reducing power, but also precursors, produced by intermediate pathways. Terpenoids represent the largest and structurally most diverse class of plant metabolites and are major products of GT biosynthesis. Despite their multiplicity, with over 30 000 well-known structures, they are all assemblies of C5 isoprene units built from isopentenyl diphosphate (IPP) and its isomer dimethylallyl diphosphate (DMAPP). There are two identified pathways for IPP and DMAPP production: i) the plastidial 2-C-methyl-D-erythritol 4-phosphate (MEP) pathway from pyruvate and glyceraldehyde-3-phosphate or ii) the cytosolic mevalonate (MVA) pathway from acetyl-CoA [6]. Although these pathways are thought to be largely independent, some exchange of precursors may occur [7], and such cross-talk requires further investigation. For instance, is there some cross-talk of plastidial and cytosolic pathways providing the 5-carbon precursors, as suggested in the latest work in peppermint [8]? And if so, what effect does it have on overall productivity? Beyond this, a major issue is the source of energy and its distribution to understand how GTs achieve their high productivity. The question becomes more intriguing when one realises that some of the GTs contain photosynthetically active chloroplasts (as the type VI GT in *S. lycopersicum* [9]). Considering that in the case of many plants where the seeds are green during embryogenesis, the light can influence the fatty acid synthesis, and potentially power refixation of CO_2_ [10, 11], we took the challenge to understand whether a similar **effect of light** can be predicted in trichomes. Till now it is still unclear what the advantages and disadvantages of photosynthetic GTs are in contrast to non-photosynthetic GTs. Moreover, there is limited research on GTs ability to absorb light. Even research focused on light intensity-mediated trichome production does not focus on their photosynthetic activity [12]. The separation of cytosolic and chloroplast-bound pathways, as well as the utility of photosynthesis, are until now only vaguely understood, and the most recent summary of current advances has been recently provided [13].

To shed light on the advantages of photosynthetic GT for terpenoid synthesis and secondary metabolism, investigations of the system’s bioenergetics and reaction flux distributions are needed. Mathematical, computational models provide a coherent framework to study metabolism. Constraint-based stoichiometric models [14] are particularly adequate for exploratory studies of the systemic properties of a metabolic network and investigations of the flux distributions. Such models are static and represent mathematically the network of biochemical reactions of an organism in the form of a matrix [15]. They can focus on various scales, with genome-scale metabolic models (GEMs) aiming at representing the whole biochemical network of an individual organism. GEMs are constructed by assigning biochemical functions to enzymes encoded in the genome, and due to the expansion of the whole genome sequencing, many plant GEMs are currently available, with *Oryza sativa indica* [16], *Arabidopsis thaliana* [17] and *Solanum lycopersicum L*. [18] among many others. Flux Balance Analysis (FBA) [19, 20], a mathematical method that allows calculating the flow of metabolites through the network, is a popular tool to predict the production rate of the compound of interest. FBA requires two assumptions: i) the experimental system is at a steady state, and ii) the network is optimised to maximise or minimise certain biological outcomes, for instance, its biomass. The so-called, cell-specific, objective functions in GEMs are optimised in a linear programming approach in which all reaction fluxes are constrained within given boundaries. This constraint-based analysis of GEMs allows the calculation of optimal flux solutions in different conditions, therefore allowing investigations on the metabolic fluxes and bioenergetics of systems.

In this work, we have reconstructed the metabolism in the photosynthetic glandular trichome type VI of a *Solanum lycopersicum* LA4024 using previously published transcriptome and metabolome data [5]. With a general, mathematical framework, we investigated the effect of having photosynthetically active machinery inside of a trichome and systematically tested the model on how GTs achieve high metabolic productivity proposed by Balcke *et al*. [5]. In our simulations, we observed the increase in terpenoid production under increasing light intensities. Increased photosynthetic activity shifts the partitioning of uptaken carbons from catabolism to anabolism due to increased energy levels. Bioenergetics and energy levels determine which of the known terpenoid precursor production pathways (MEV, MEP) is more desirable/optimal in different light/stress conditions. Our model can explain the benefits of having chloroplasts in GTs and serves as a groundwork for further investigations of the possible cross-talks between the two pathways of terpenoid precursor synthesis. It complements the previous work by Balcke *et al*. [5] by not only confirming their hypothesis that the light-dependent reactions of photosynthesis support the secondary metabolite pathways, but explaining how this support is achieved. Finally, our predictions have been tested *in vivo* and we provide an experimental validation by showing that under high light conditions, production of the most abundant MEP-derived terpenes (2-carene and *β* -phellandrene/D-limonene) increases, whilst most abundant MEV-derived terpenes (*β* -caryophyllene and *α* -humulene) remain unchanged.

## 2 METHODS

### 2.1 Choice of the model organism

In this study, we have chosen to investigate type VI GT in the tomato genus. The tomato genus displays seven types of trichomes: II, III and V (non-glandular) and I, IV, VI and VII (glandular trichomes), with type VI being the most abundant one in the *Solanum lycopersicum* species. *S. lycopersicum* serves as an excellent model organism for glandular trichome study due to the availability of i) high-quality complete genome sequence [21], ii) excellent genetic resources [22], iii) comparative multi-omics data [5], iv) several mathematical models available, including whole genome metabolic network reconstruction [18], and v) in contrast to other well-studied organisms like peppermint [23, 24], possession of photosynthetic GT.

### 2.2 Modelling environment

Our model is implemented in Python, using our in-house developed package moped, “an integrative hub for reproducible construction, modification, curation and analysis of metabolic models” [25]. With moped all decision processes and modelling steps are well documented in a transparent and repeatable fashion. All details and information about the exact construction process of the model, as well as all investigations and analyses, can be found in our provided scripts at https://gitlab.com/qtb-hhu/models/glandular-trichomes. The summary of the construction steps is provided in the section below.

### 2.3 Model construction and assumptions

Although a genome-scale model of tomato metabolism is available (iHY3410 model [18]), we decided to use a bottom-up approach and perform the reconstruction ourselves, because we were not able to reconstruct the steps of manual curation performed by the authors. We based the model reconstruction on available transcriptomics and metabolomics data [5], the LycoCyc database (tomato metabolic pathway database, version 3.3 [26], available from Solanaceae Genomics Network, http://www.sgn.cornell.edu) and biochemical knowledge in plants from scientific publications. All reactions and metabolites found in transcriptomics and metabolomics data have been added to the model from the LycoCyc database using the moped metabolic modelling package, ensuring GPR rules for all added reactions in the network.

We used Meneco, a tool for metabolic network completion [27] to subsequently fill gaps in our network with annotated reactions from the LycoCyc database, so our model is capable of synthesizing all compounds found within the metabolomics data [5], all terpenoids found in photosynthetic GTs of tomato [28] as well as all amino acids, nucleotide bases and lipid precursors from sucrose, light, orthophosphate, ammonia, sulfate, protons and water. All reactions in our model have been checked for mass and charge balance and are able to carry steady-state fluxes. Our model shows the ability to synthesize biomass precursors and terpenoids on a realistic scale. The model has been examined for inconsistencies in energy metabolism by analyzing the model behaviour according to changes in ATP demand. Increasing ATP demand leads to plausible changes in key reactions of the model, such as a decrease in objective function flux. A detailed description of every implemented step in model construction, as well as the code for reproducing the entire model reconstruction process, can be found in our *model_construction*.*ipynb* notebook in the supplementary material.

Our model is a data-driven, yet simplified, constraint-based model which is ensured not to include infeasible energy and mass-generating cycles. Within our model simplifications, we found that a model consisting of three essential compartments (cytosol, intermembrane space and extracellular space, as represented on Fig. 1) is able to represent photosynthetic GT metabolic profiles. While detailed compartmental separation is common practice in large genome-scale metabolic models, it would not make any difference to the results of our model simulations due to the fact that there are several intercompartmental transporters between the chloroplast and the cytosol for energy equivalents like ATP and other key metabolites [29]. Adding over-detailed compartmentalisation to the model would therefore not alter any of our results and is left out for the sake of model simplicity and preventing unfavourable model modifications. The resulting model consists of 1307 reactions and 1371 metabolites and thanks to the integration of the multiomics data, its behaviour has been ensured to match reported experimental observations [5]. There are nine exchange reactions, allowing the free exchange of inorganic metabolites such as oxygen, as well as light absorption and sucrose uptake. To ensure the highest quality and consistency our model has been thoroughly inspected using the MEMOTE standarised testing suite [30].

**FIGURE 1.**
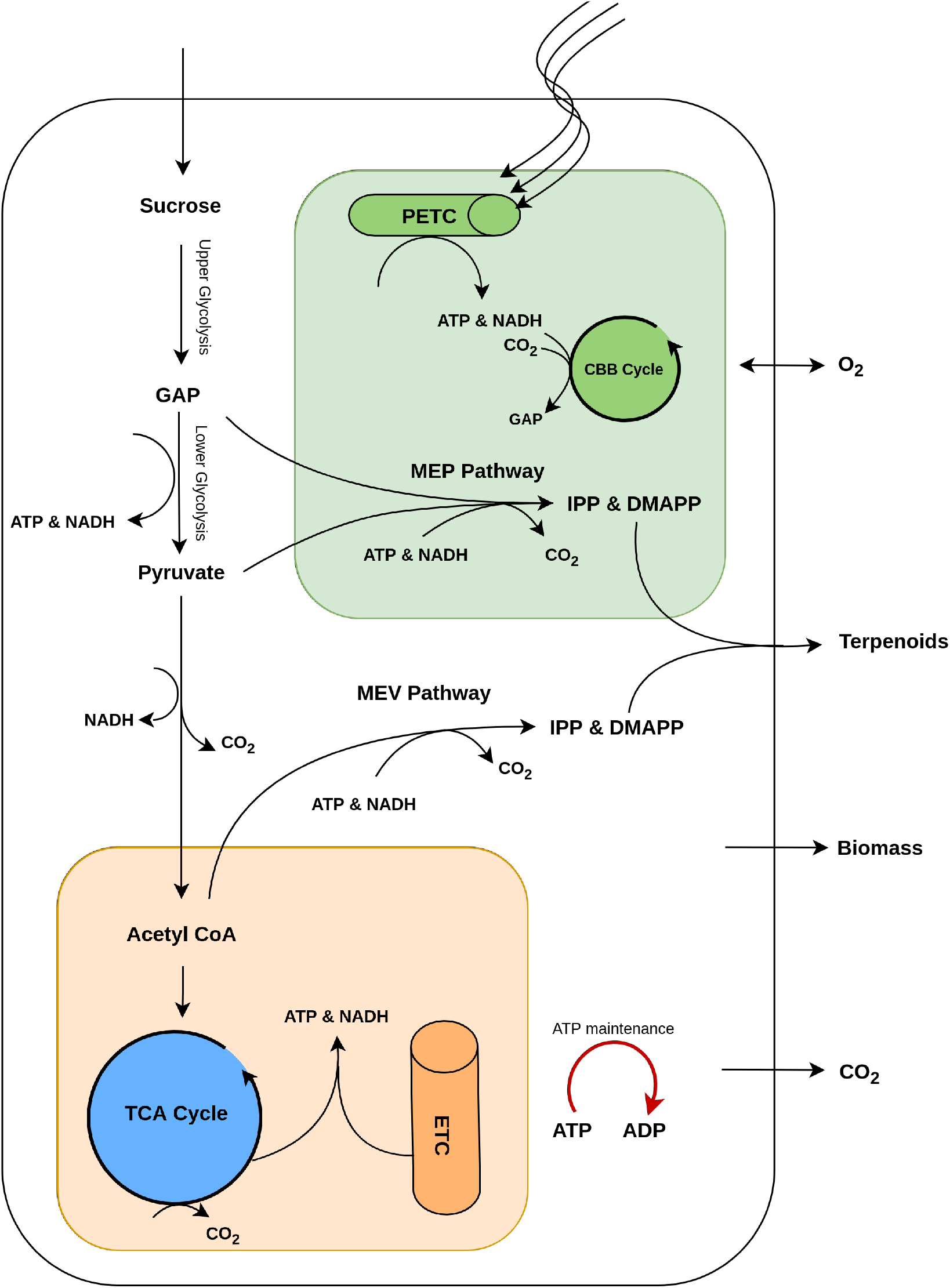
Schematic overview of the key processes included in a constraint-based model of photosynthetic glandular trichome (GT) metabolism. While the model is built using transcriptome and metabolome data and includes a large number of reactions, only pathways and metabolites of importance to the results are highlighted in the presented model scheme. These include CBB Cycle, Calvin-Benson-Bassham Cycle; DMAPP, dimethylallyl diphosphate; IPP, isopentenyl diphosphate; MEP, methyl-erythritolphosphate pathwy; MEV, mevalonate pathway; PETC, photosynthetic electron transfer chain; TCA Cycle, tricarboxylic acid Cycle.

#### 2.3.1 Optimisation and objective function

In most constraint-based models and their analyses, the maximization objective is the production of biomass [31]. While this may be applicable for prokaryotic organisms, we doubt that photosynthetic glandular trichome cells are maximising the increase of their replication rate, and rather maximise terpenoid synthesis while also having a mandatory production rate of macromolecules to keep cells intact. For this, our model includes an objective function to produce terpenoids while requiring a fixed flux through a function of biomass synthesis, consuming typical components like amino acids, sugars, nucleotides, cell wall, and fatty acid precursors. We used a simplified biomass function inspired by plant biomass functions from Seaver *et al*. [32], similar to standard biomass functions used succesfully for FBA in plants [33, 34]. Our aim was to capture the necessity for growth and self-repair, while setting the objective function to maximize the production of terpenoids. To describe additional energy required for the maintenance of cells, we implemented a representative reaction for ATP maintenance, as it is common practice in metabolic modelling [35].

#### 2.3.2 Calculation of flux units and light intensity units

There is limited research on GTs ability to absorb light (e.g., [36]), and we have not found any dedicated research focused on the optical properties of GTs. Overall, while there is some evidence to suggest that GTs may have optical properties allowing for some light absorption, more research is needed to fully understand their ability to absorb light. Therefore we have decided to estimate the maximal absorption rate based on the reported maximal production fluxes. Light is therefore represented as **photons absorbed by the photosystems** used for photosynthesis in contrast to the incident light that will be several-fold higher.

Although it is known that due to diel cycles of photosynthesis different metabolic flux patterns in the light and the dark are observed and require different treatments for the optimisation problem[37], our FBA on the trichome model is performed under continuous light. The units of light absorption are represented in 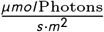 and the detailed calculations are provided in the Supporting Information. As it has been reported that carbon dioxide exchange is 100 times lower in photosynthetic GTs than in leaves we decided not to include a carbon dioxide influx, however, carbon dioxide is produced in the system and can flow out of the model [5].

A suggested terpenoid production rate of GTs has been provided by Turner *et al*. [38] at 0.0017 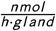. Assuming that this rate can be applied to the maximal terpenoid production rate of photosynthetic GTs of tomatoes, we transform our calculated fluxes to the corresponding units by 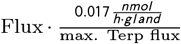. Next, in order to convert the fluxes of photons into units of light intensities, we calculated the light-absorbing surface of the GT type VI. Although the head of GT type VI is made up of 4 secretory cells, we simplify the whole surface of the head as a sphere. Based on the measured values of the diameters of GTs from [39] and bright-field microscopic image from [9] we took the estimate of 50 *μ*m as the diameter. This number can be substituted with a different value and the light conversion function will be adapted. Under these assumptions, the surface area can be estimated as:

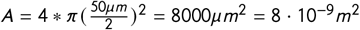

To calculate the conversion factor for the photon absorption of GTs, we first calculate the units of photons absorbed by the gland at saturated light flux and maximal terpenoid production predicted by our model as:

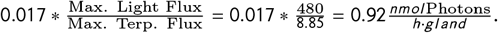

To convert this unit into 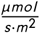, we first calculate the corresponding unit for 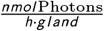 by:

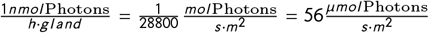

for our maximal Light flux, this corresponds to 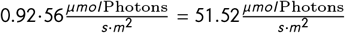 as the saturating light intensity, providing the light flux conversion factor of Light 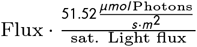.

### 2.4 Experimental set-up

The high light experiment was conducted using two LED-light panels (Rhenac GreenTech AG, Hennef, Germany) placed inside a phytochamber and separated by a black curtain. Tomato (*Solanum lycopersicum* cv Moneymaker) plants were germinated on soil under control conditions (CN): 16h light, 422 *μ* mol *m*^−2^*s* ^−1^ at 25°C and 8h dark at 20°C, 70% humidity day and night. After 22 days of growth, half of the plants were transferred to high light (HL) conditions for further 7 days, where light intensity was adjusted to 1289 *μ* mol *m*^−2^*s* ^−1^. Light spectra were recorded using a Specbos 1211UV (JETI, Jena, Germany) photometer (Fig. S1). Volatile terpenoids of leaflets of three different developmental stages were collected by surface extraction using leaf discs of 1 cm diameter and vortexed for 30 seconds in n-hexane. Mono- and sesquiterpenes were detected using a Trace GC Ultra gas chromatograph coupled with an ATAS Optic 3 injector and an ISQ mass spectrometer (Thermo Scientific) with electron impact ionization. The chromatographic separation was performed on a ZB-5ms capillary column (30 m × 0.32 mm, Phenomenex). The flow rate of helium was 1 ml min1, and the injection temperature rose from 60 to 250°C at 10°C sec^1^ during 30 sec. The GC oven temperature ramp was 50°C for 1 min, 50–150°C at 7°C min^1^ and 150–300°C at 25°C min^1^ for 2 min. Mass spectrometry was performed at 70 eV in full scan mode with *m*/*z* from 50 to 450. Data analysis was done with the Xcalibur software (Thermo Scientific).

## 3 RESULTS

We used our model to perform a general analysis in which we simulate the rate of terpenoid synthesis over systematically increasing light intensities via parsimonious Flux Balance Analysis (pFBA) [40]. Fig. 2 displays that with increasing light absorption, the rate of terpenoid synthesis in photosynthetic GTs increases up until approximately 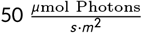. This increase in terpenoid synthesis rate with increasing absorbed light is particularly interesting due to the fact that the model can not utilise atmospheric carbon dioxide, and sucrose is the only carbon source. This means that there is a change in metabolic fluxes which enables this increase in terpenoid synthesis rate. To further investigate what changes in the metabolism of photosynthetic GTs in increasing light intensities, we inspect the respective changes in the exchange fluxes of the model. Fig. 2 shows the exchange fluxes of carbon dioxide and oxygen in our pFBA model simulations over increasing light absorptions. Noticeably, the release of carbon dioxide systematically decreases up until approximately 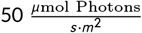. Interestingly, the consumption of oxygen decreases to zero at approximately 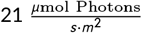. From this light intensity on, oxygen release begins and increases until 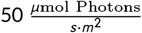. These observations are crucial for a general understanding of the model behaviour. An increase in absorbed light causes higher photosynthetic activity, resulting in oxygen production. This explains the decreasing oxygen uptake and the switch to oxygen release at 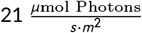 absorbed light. However, the steady decrease in carbon dioxide excretion is especially noteworthy. Most carbon dioxide is produced within catabolism, therefore the model behaviour hints at a decrease in catabolic activity in higher light intensities.

**FIGURE 2.**
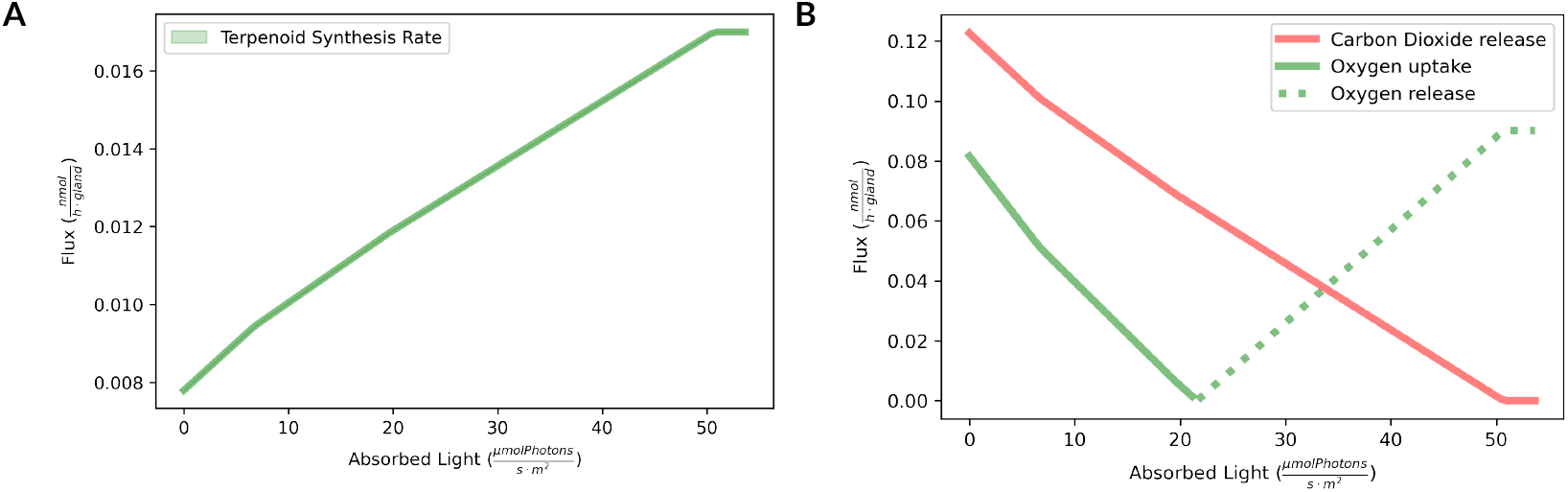
Impact of rates of absorbed light on the predicted fluxes through the photosynthetic glandular trichome. **a** Terpenoid synthesis flux over different rates of absorbed light. The rate terpenoid synthesis increases with higher amounts of absorbed light. **b** Oxygen and carbon dioxide exchange fluxes over different rates of absorbed light. Increased light absorption decreases carbon dioxide release and oxygen absorption. In higher rates of light absorption, oxygen is secreted.

To investigate how the catabolic activity in our model simulations changes over increasing light intensities, we further inspect representative reactions for relevant catabolic pathways in our model. As sucrose, a disaccharide is the only carbon source in our model, we inspect representatives of the upper glycolysis, the lower glycolysis and the TCA cycle. Fig. 3 displays the fluxes of these reactions over different light intensities (shown on the x-axis as fractions of saturating light intensities) relative to their fluxes in the dark. The sucrose synthase and saccharase represent upper glycolysis activity. The 6-phosphofructokinase, GAP dehydrogenase, and pyruvate kinase represent lower glycolysis activity and the pyruvate dehydrogenase and the citrate synthase represent TCA cycle activity. Furthermore, the RuBisCO rate is displayed to monitor the rate of carbon refixation. The results show that fluxes of upper glycolysis remain completely unchanged in increasing light intensities, however, the fluxes in lower glycolysis decrease in higher light conditions. An even higher impact can be observed for the TCA cycle activity. The pyruvate dehydrogenase activity steadily decreases, and the citrate synthase activity abruptly decreases in increasing light conditions. These observations show that catabolic pathways which are not responsible for energy and redox equivalent production (like upper glycolysis) are unaffected by increasing light intensities. However, the lower glycolysis and the TCA cycle, both catabolic pathways that produce energy and redox equivalents, display a strong flux decrease in higher light conditions. There is no reason to think that trichomes would not have the same regulatory mechanisms as mesophyll cells, leading to decreased glycolysis in higher light conditions. Increased photosynthesis is accompanied by increased photorespiration, which eventually supplies reducing equivalents in the mitochondria, which can then be used to fuel the respiratory electron transport chain [41]. The increase in terpenoid synthesis flux observed in Fig. 2, and the decrease in catabolic fluxes in Fig. 3 strongly suggest that increasing light conditions **shift the carbon partitioning from catabolic to anabolic pathways**. This shift is enabled due to the energy and redox equivalent production of the photosynthetic electron transport chain in photosynthetic GTs. The metabolic network is not dependent on the energy from oxidising carbon bodies in high-light conditions, and can therefore use more of those carbon bodies in terpenoid synthesis pathways. Interestingly, RuBisCO activity increases in higher light intensities, displaying that only very high levels of photosynthetic energy supply allows the refixation of carbon that is lost as carbon dioxide in anabolic processes (like terpenoid synthesis). GAP, pyruvate and acetyl-CoA are carbon bodies which can be used to produce either energy and redox equivalents or terpenoid precursors. Acetyl-CoA is the initial substrate of the TCA cycle in which it is oxidised to gain energy and redox equivalents but is also the initial substrate of the MEV pathway, also known as the isoprenoid pathway, which is the primary terpenoid synthesis pathway in non-photosynthetic GTs. GAP and pyruvate are metabolites within the lower glycolysis pathway and also initial substrates of the MEP which is a terpenoid synthesis pathway present in photosynthetic GTs.

**FIGURE 3.**
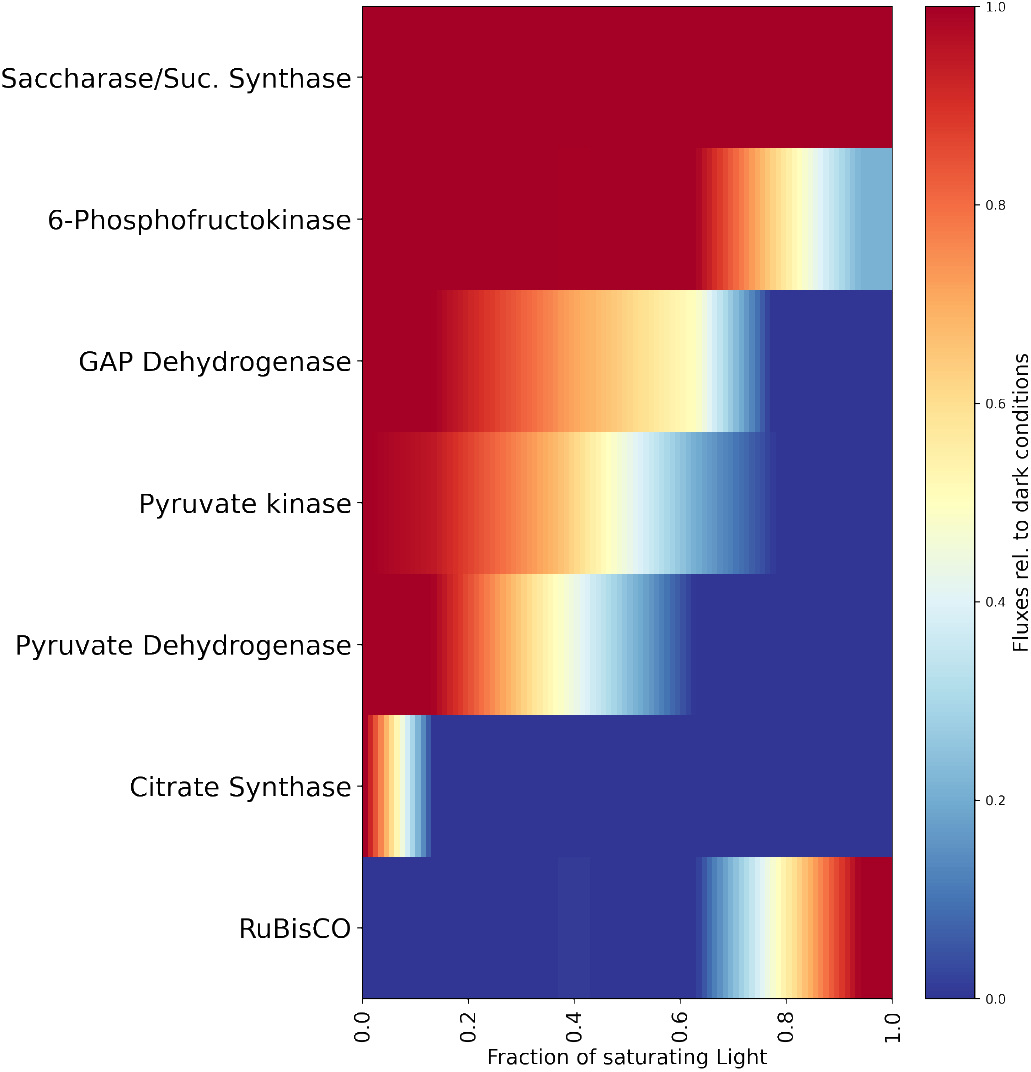
The relative fluxes of six selected catabolic reactions and one carbon fixation reaction calculated for increasing fractions of saturating light. The fluxes are normalised to the respective fluxes under completely dark conditions. Reactions of upper glycolysis remain unaffected by increased light absorption, while the fluxes of reactions of lower glycolysis and the TCA cycle decrease with increased absorbed light. Additionally, RuBisCO flux is only present when light absorption is high.

To further analyse how the consumption of these metabolites depends on the illumination, we simulated the relative consumption rate of GAP/pyruvate and Acetyl-CoA by the aforementioned pathways over increasing light intensities. Fig. 4 displays the proportions of the consumption of these compounds by the TCA, MEV and MEP pathways. In low light intensities, more than half of the substrates are consumed by the MEV pathway, and the remainder is consumed by the TCA cycle, in both cases in the form of acetyl-CoA. In higher light intensities, the fraction of substrates consumed by the TCA cycle is decreasing until it does not consume any more substrates. At this point, the relative flux of lower glycolysis starts decreasing, and the MEP pathway is beginning to consume proportions of the substrates, gradually taking over. This is a very important observation that shows that increasing light intensities, leading to higher energy levels due to photosynthetic activity, shift the carbon partitioning from catabolic to anabolic pathways by reducing the TCA cycle and lower glycolytic flux and increasing terpenoid synthesis. Furthermore, it shows that the two terpenoid synthesis pathways, MEP and MEV, are more advantageous at different energetic levels. In lower light intensities, and therefore lower energetic levels, the MEV pathway seems to be more advantageous because the conversion of GAP and pyruvate to acetyl-CoA produces energy and redox equivalents, and the resulting acetyl-CoA can directly be used in the TCA cycle to generate additional energy and redox equivalents. In higher light intensities, and therefore higher energetic levels, the MEP pathway is more advantageous because the high energy levels provided by photosynthetic activity remove the necessity of providing energy and redox equivalents via lower glycolysis and the TCA cycle. Instead, GAP and pyruvate can directly be used as substrates with higher energy contents (than acetyl-CoA) in the MEP pathway, and therefore further increase the fraction of carbon used in anabolism, enabling more efficient terpenoid synthesis.

**FIGURE 4.**
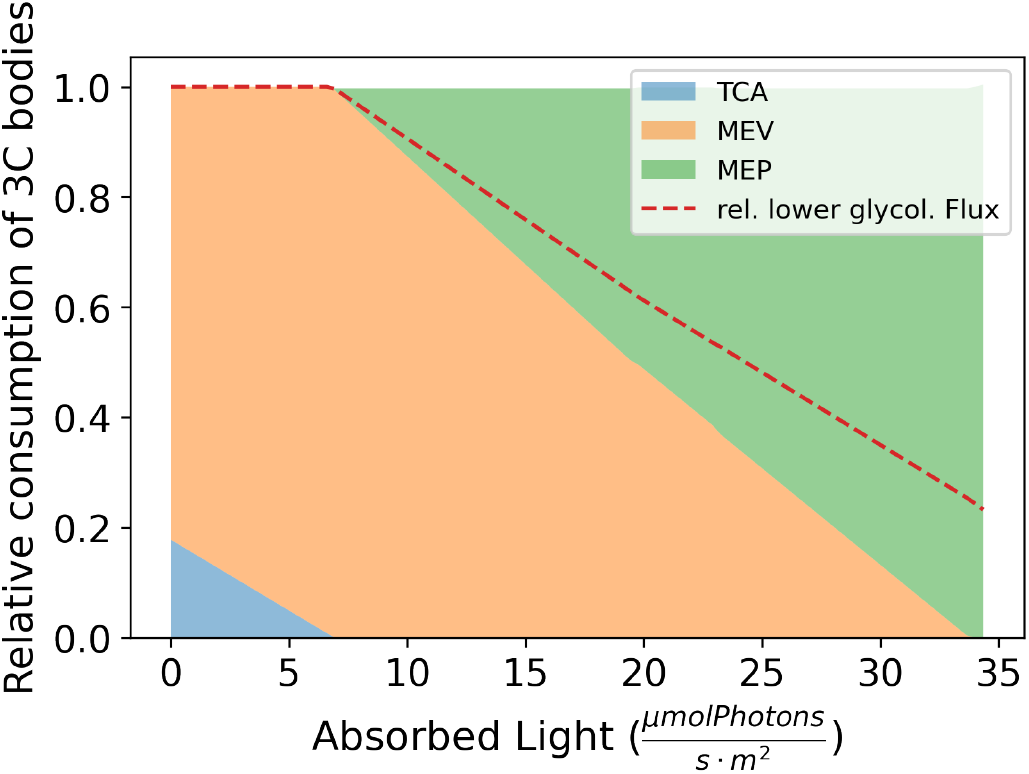
Predicted relative consumption of GAP, pyruvate and acetyl-CoA (here described as 3C bodies) by different pathways over increasing fractions of saturating light. The fraction of the lower glycolysis flux relative to dark conditions is displayed as a dashed line. Increasing rates of absorbed light decrease the fraction of 3C bodies consumed by catabolic pathways, like lower glycolysis and TCA cycle. Furthermore, the fraction of 3C bodies consumed for terpenoid synthesis switches from the MEV to MEP pathway in high rates of light absorption.

This phenomenon can also be observed in Fig. 6, in which we used model simulations to calculate the fluxes of the final MEV and MEP reactions in systematically changing light conditions and ATP maintenance costs. In this analysis, higher ATP maintenance costs reflect increased energy requirements of cells in e.g. stress conditions. At low light conditions and low ATP maintenance costs, the MEV pathway is the main terpenoid synthesis pathway, with very little MEP pathway activity. In low light conditions and high ATP maintenance costs, the MEV pathway is the only active pathway. However, the overall terpenoid synthesis flux is relatively low due to the increased demand for catabolic flux in such conditions. At high light conditions and high ATP maintenance costs, the MEP pathway is carrying the majority of terpenoid synthesis flux. In high light conditions and low ATP maintenance costs, the MEP pathway is the only active terpenoid synthesis pathway, providing the highest terpenoid synthesis flux. It appears that the distribution of terpenoid synthesis between the MEV and the MEP pathways is highly dependent on the light conditions and resulting energy levels of the photosynthetic GTs.

The general conclusions of our model simulations have then been tested experimentally. The impact of light intensity on the shift from the MEV to MEP precursor pathway has been tested by quantifying sesquiterpenes, produced by MEV, and monoterpenes, produced by MEP. Three-week-old tomato (*Solanum lycopersicum* cv Moneymaker) plants were exposed to nearly threefold higher light intensity (HL) for seven days and the productivity of the main volatile terpenoids produced in the type VI GTs was estimated by GC-MS along different leaf developmental stages. The results showed a significant increase in monoterpenoids while the sesquiterpenes remained unchanged and such increment is linked to the age of the leaves (Fig. 5). These findings suggest that the photosynthetic light reactions support the productivity of the specialised reactions occurring in the plastids through the MEP pathway.

**FIGURE 5.**
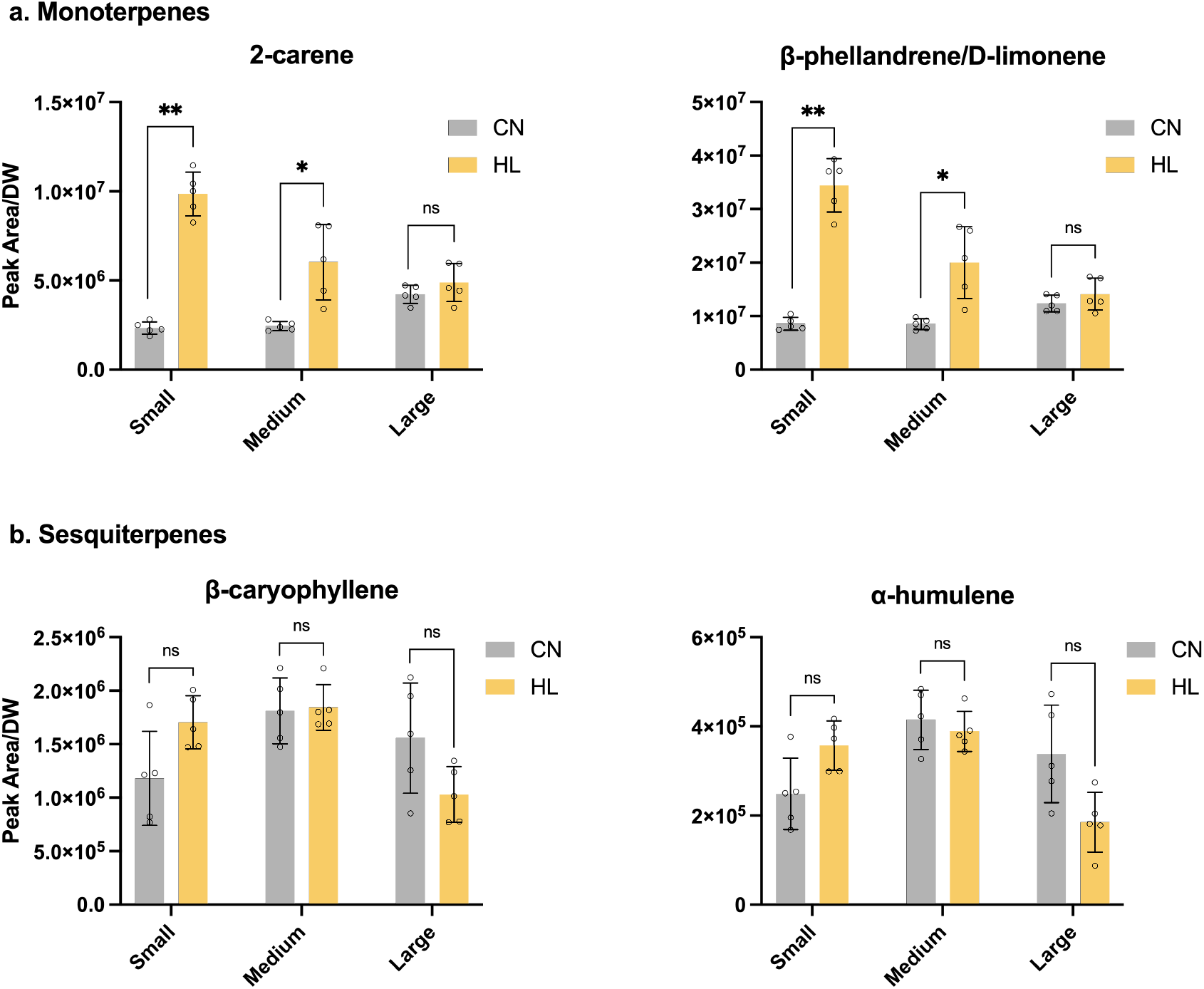
Effect of high light in volatile terpene content of leaves of tomato. Estimation of (a) monoterpenes and (b) sesquiterpenes by gas chromatography-mass spectrometry (GC-MS) on young (Small, from bottom-to-top leaf number 6), expanded (Medium, leaf number 5) and fully develop (Large, leaf number 4) leaves, in control (CN) and high light (HL) conditions. Chromatogram peak areas were normalized by leaf dry weight (DW). Error bars indicate SD (*n* = 5 biological replicates; ^*^P < 0,05, ^**^P < 0,01; using *t*-test).

Finally, the high rate of terpenoid synthesis in high-light conditions is partly resulting from increased rates of carbon refixation. It remains unknown how active the CBB cycle is in photosynthetic GTs. To quantify the impact of different carbon refixation fluxes, we performed a systematic analysis in which we calculated the terpenoid synthesis rate over different quanta of **absorbed light** and systematically changed the activities of RuBisCO (Fig. 7). Interestingly, the overall rate of carbon refixation is increasing the rate of terpenoid synthesis by almost 20%, while the shift in carbon partitioning between catabolism and anabolism increases it by almost 200%. This shows that the impact of energy-dependent shift in carbon partitioning and isoprenoid synthesis pathways is a lot higher than the RuBisCO-dependent refixation of carbon dioxide.

**FIGURE 6.**
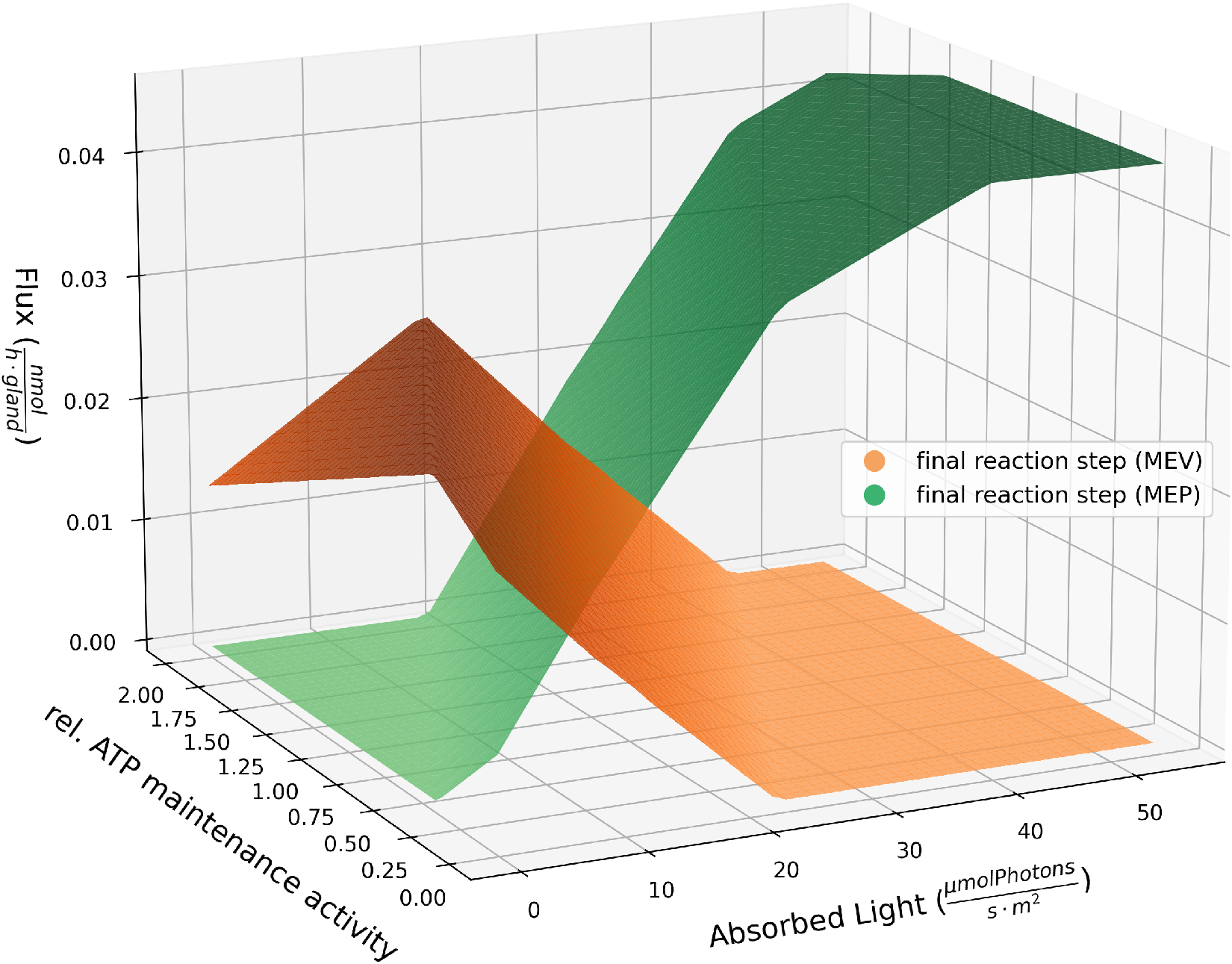
Fluxes of the final reaction steps of the MEV and MEP pathway over increasing relative ATP maintenance activities, as well as increasing rates of light absorption. In energetically favorable conditions, like high light and low ATP maintenance (function representing the additional energy requirement for the maintenance of cells), terpenoid synthesis is carried out by the MEP pathway. In opposite conditions, meaning low light and high ATP maintenance, the MEV pathway is performing terpenoid synthesis.

**FIGURE 7.**
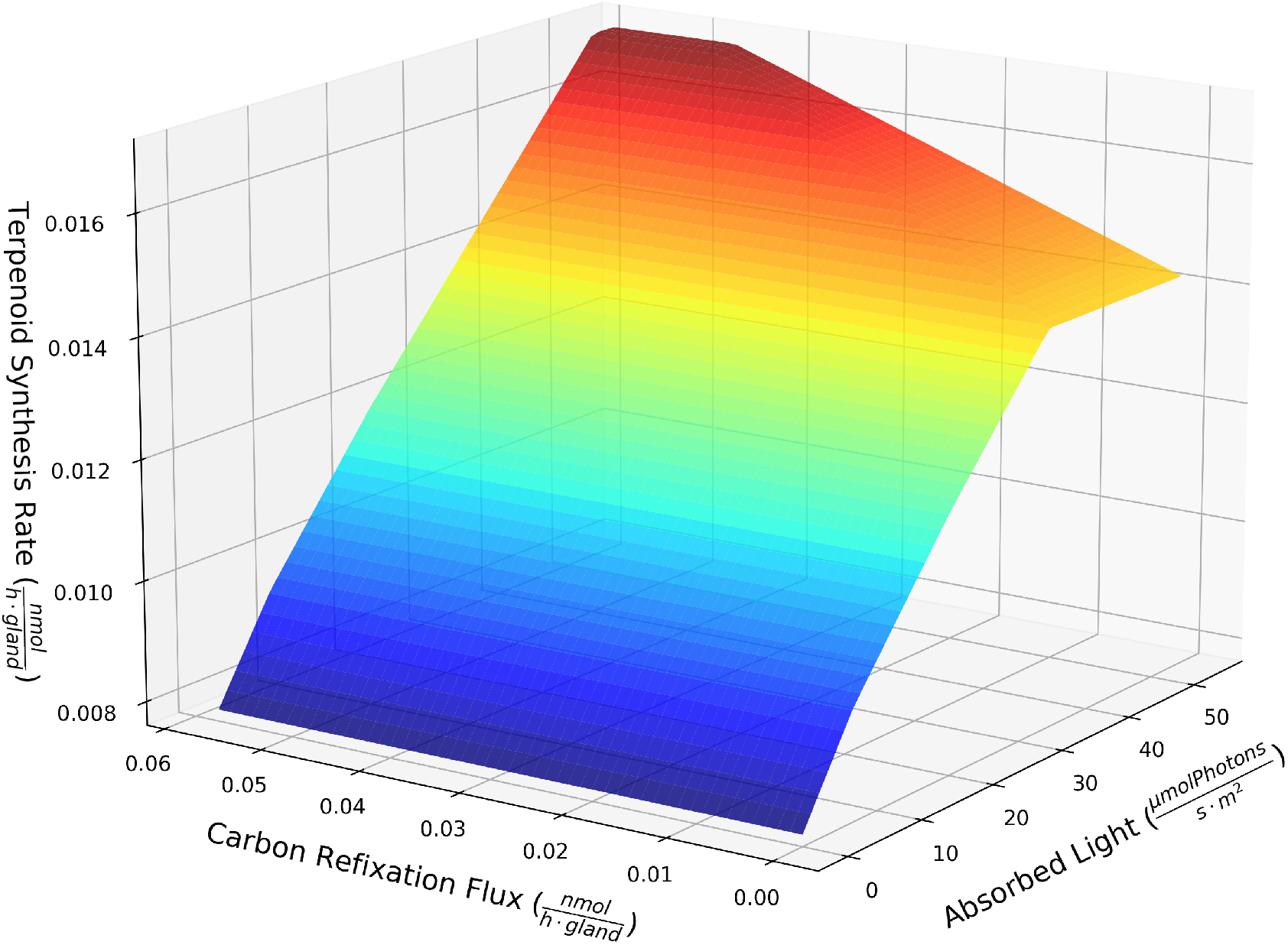
Predicted flux of the terpenoid synthesis under changing carbon refixation rates, as well as increasing rates of light absorption. Under simulated saturating light, the rate of terpenoid synthesis is two-fold higher than in darkness. If additionally the carbon refixation flux is increased to the maximal chosen value, the increase in terpenoid synthesis is less than 20%. This result shows how the energy-dependent shift in carbon partitioning and isoprenoid synthesis pathway is higher than the RuBisCO dependent refixation of carbon dioxide.

## 4 DISCUSSION

In photosynthetic GTs, synthetic pathways of terpenoids and other secondary metabolites are found in the cytosol of the cells and the chloroplasts. The additional terpenoid synthesis pathway in photosynthetic GTs has been subject to many speculations, e.g., terpenoid production in chloroplasts is specialised for the production of particular secondary metabolites [28]. In our work, we built a simplified, yet data-driven constraint-based model of photosynthetic glandular trichome metabolism, and used it to show that one of the two different synthesis pathways is more advantageous for terpenoid production than the other in different energy availabilities. Previously published multi-omics data [5] supports our hypothesis that the energy and reducing power (ATP and NADPH) from photosynthesis are primarily used to power the secondary metabolism. Our model provides a mechanistic explanation of how this is achieved.

We show that with lower energy availability, the cytosolic MEV pathway is more advantageous for terpenoid synthesis because the catabolic pathways, producing the critical initial substrate acetyl-CoA from sucrose, provide additional energy and redox equivalents needed for all cellular activities, including terpenoid synthesis. However, higher energy availability (coming from photosynthetic activity in higher light conditions) removes the need for the additional energy and redox equivalents gained from the conversion of sucrose to acetyl-CoA. Therefore substrates with higher energy levels (GAP and pyruvate) can be directly used for terpenoid synthesis. This shortcut of catabolic reactions reduces the loss of carbon as carbon dioxide and increases the flux of carbon through anabolic processes. The general conclusions derived from the theoretical analyses have been strengthened with the experimental evidence that under higher light intensities production of monoterpenoids is significantly increased, whilst the production of sesquiterpenes remains unchanged (Fig. 5).

We show that in higher light conditions, energy levels of the photosynthetic GTs are so advantageous, that excess energy can be spent to perform carbon refixation using the CBB cycle. In the supplementary material, we calculated that the terpenoid yield per sucrose is twice as high in high light compared to low light conditions. This illustrates that the benefit of including chloroplasts in GTs is not only the ability to shift carbon partitioning from catabolic to anabolic processes but also to further maximize carbon use efficiency. It is important to note that the increase in terpenoid synthesis from carbon refixation is not nearly as high as the increase from the shift in carbon partitioning, as seen in Fig. 7. This dual behaviour, i.e. in the absence versus in presence of photosynthesis, is reminiscent of the situation in photosynthetic leaves, where the TCA cycle is inactive during the day and active during the night [42].

Interestingly our model shows that even without CBB cycle activity, the TCA cycle may be reversed in high energy availability and function as a reductive TCA cycle. This reductive TCA cycle could theoretically take over the function of the CBB cycle, using energy to fix carbon dioxide which was produced in catabolic and anabolic reactions, thus increasing carbon use efficiency. This is a very interesting observation, as, from a bioenergetic point of view, such a scenario is possible. Considering the previous results where Phosphoenolpyruvate carboxylase (PEPC) expression was significantly increased in trichomes compared to leaves [5], we consider PEPC as a plausible candidate to mediate the carbon fixation. However, we decided to adjust the key reactions of the TCA cycle for this scenario as irreversible to prevent this phenomenon to be included in our results for now. The reason for this decision is that the reductive TCA cycle is usually found in green sulfur bacteria and different thermophilic prokaryotes and archaea [43, 44]. This indicates that from a phylogenetic perspective, the presence of a reductive TCA cycle in photosynthetic GTs is rather unlikely. However, we think that this model suggestion is worth investigating the fluxes of the TCA cycle in light conditions in photosynthetic GTs, as it has been suggested that carbon dioxide may be recovered [45]. Generally, instead of showing that chloroplastic terpenoid synthesis pathways provide improved production of particular terpenoids, our work shows that the chloroplast in photosynthetic GTs functions as a solar panel in light conditions, which can be used to shift carbon from catabolic to anabolic fluxes and even enable carbon dioxide refixation and therefore improve carbon use efficiency. To support our findings, experiments are needed which can keep track of the rate of terpenoid synthesis in similar sucrose availability but different light absorptions. Interestingly, such an increase in the efficiency of carbon use through RubisCO, but without the CBB cycle, has been observed in other plant cells. Schwender *et al*. [46] showed that Rubisco without the Calvin cycle improves the carbon efficiency of developing embryos of *Brassica napus L*. (oilseed rape) during the formation of oil.

Photosynthetic carbon refixation indicates that photorespiration may be present in photosynthetic GTs. Although photorespiratory genes were very low expressed in the transcriptome data [5], and photorespiration is not included in our model, we show that in high light there is oxygen evolution in photosynthetic GTs. Therefore, new experimental data obtained in high light intensities and gas exchange rates is required to investigate putative photorespiratory activities. Furthermore, it remains unclear if and how high the evolution of reactive oxygen species and photodamage is present in photosynthetic GTs. For this, quantitative metabolic data for the components of the electron transport chain is needed, as well as measurements of the photosynthetic efficiency in photosynthetic GTs. Finally, more questions regarding the dynamics, and not only bioenergetics of trichomes arise. E.g., what is the composition of terpenoids under different light intensities, or even light colours? A recent study using different basil cultivars showed that light spectra affect the concentrations and volatile emissions of important compounds [47]. Such light modulation requires further investigation with a use of more detailed models of secondary metabolism in photosynthetic GTs that take the light spectrum into consideration. For transparency, we include the light spectra used for our experimental validation in Fig. S1. As most of the processes discussed here are heavily dependent on enzyme kinetics and saturation, constraint-based models like the one presented may not be the best method for answering these new emerging questions. Mechanistic models, e.g. based on ordinary differential equations, can include such information (if available) and may be helpful to give further insights into terpenoid synthesis in photosynthetic GTs. Further interdisciplinary studies combining experiments and robust theoretical simulations can provide a quantitative understanding of whether, how, and by how much the production of specific terpenoids could be increased, and with this work, we provide the stepping stone for such analyses. The outcomes of this research provide quantitative measures to assess the beneficial role of chloroplast in GTs and can further guide the design of new experiments aiming at enhanced terpenoid production.

## Abbreviations

CBB: Calvin-Benson-Bassham cycle
DMAPP: dimethylallyl diphosphate
FBA: flux-balance analysis
F6P: Fructose-6-Phosphate
GEM: genome-scale metabolic models
GT: glandular trichome
IPP: isopentenyl diphosphate
LP: linear programming
MEP: methyl-erythritolphosphate pathwy
MEV: mevalonate pathway
PEPC: Phosphoenolpyruvate carboxylase
PETC: photosynthetic electron transfer chain
TCA: tricarboxylic acid

## Supporting Information

The computational model presented here, together with the Jupyter Notebook containing step-by-step instructions on how to reconstruct the model and reproduce all analyses included in this manuscript are openly available on the GitLab repository https://gitlab.com/qtb-hhu/models/glandular-trichomes or can be requested from the authors.

---

**FIGURE S1.**
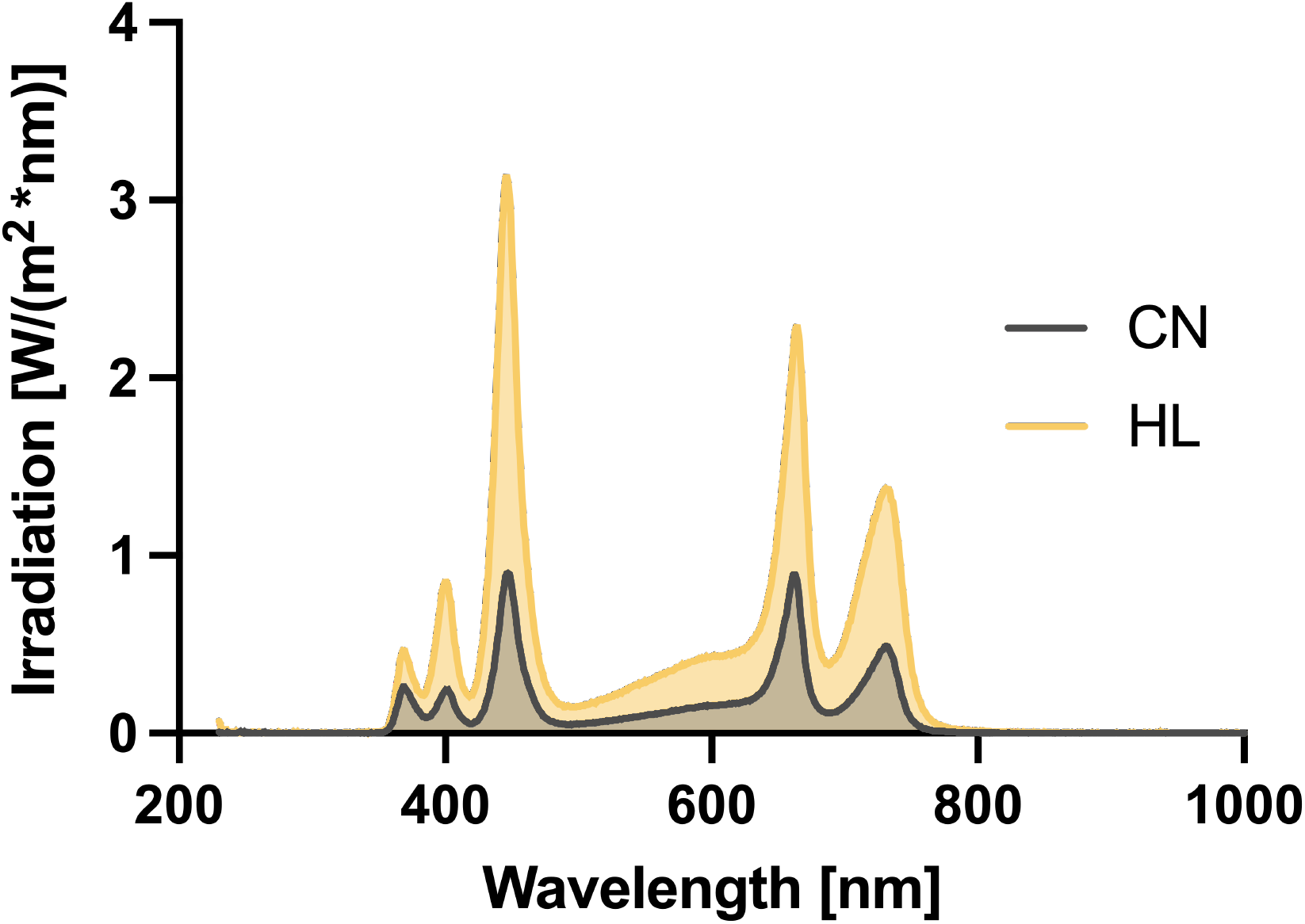
Light spectra of the two light conditions used in the experiment. Wavelength recorded from 230 to 1000 nm. CN, control; HL, high light.

